# Inhibition of serine and arginine-rich splicing factor 3 induces epidermal differentiation and decreases cutaneous squamous cell carcinoma risk

**DOI:** 10.1101/2024.11.07.622551

**Authors:** Isoline M. Donohue, Haiyin He, Audrey Nguyen, Gavin Hui, Jananee Muralidharan, William C. Pike, Michael L. Jackson, Christine Ko, Ankit Srivastava, Carolyn S. Lee

## Abstract

Serine/Arginine-rich splicing factor 3 (SRSF3) is one of 12 SRSFs that regulate gene expression via alternative splicing. SRSF3 is upregulated in cutaneous squamous cell carcinoma (cSCC) and several squamous cancer cell lines in relation to normal keratinocytes. We suppressed SRSF3 with a specific inhibitor, SFI003, and observed an increase in epidermal differentiation. Our data suggests that in cSCC, SRSF3 overexpression suppresses cellular differentiation to enable cancer progression. In a clinical setting, patients taking known SRSF3 inhibitors digoxin and amiodarone exhibited higher cSCC-free survival compared to a propensity score-matched cohort treated with beta blockers. Thus, SRSF3 upregulation may be a novel therapeutic target in cSCC that can improve patient prognoses.

## Introduction

Serine/Arginine-rich splicing factors (SRSFs) are RNA-binding proteins (RBPs) that play a crucial role in RNA processing via alternative splicing, genome stability, and mRNA transport^1^. These RBPs are localized in the nucleus but can also be found in the cytoplasm to regulate mRNA export^1,2^. SRSFs are involved in the pathogenesis of various diseases and are protooncogenic in cancers^1^. SRSF3 overexpression is associated with the progression of colorectal, ovarian, cervical, lung, liver, bladder, thyroid, and kidney cancers^2-4^. However, SRSF3 expression in keratinocyte cancers remains uncharacterized and the role of this protein in the skin is unknown. In this study, we examined the function of SRSF3 in regulating epidermal differentiation – a key process disrupted in cutaneous squamous cell carcinoma (cSCC).

## Materials and Methods

### Cell culture and treatments

Primary human neonatal keratinocytes were isolated as previously described^5^ and cultured in Medium 154 with Human Keratinocyte Growth Supplement and Penicillin-Streptomycin (ThermoFisher Scientific, USA). CAL27 cells were cultured in DMEM medium (Gibco) supplemented with 10% FBS (MilliporeSigma). SCC13 cells were cultured in Keratinocyte SFM medium supplemented with 0.3 mM CaCl_2_, 50 µg/ml of Bovine Pituitary Extract, and 5 µg/µL human EGF (ThermoFisher Scientific). A431 and SCCIC1 cells were cultured as previously described^5^. SFI003 (TargetMol Chemicals Inc. supplied by Fisher Scientific, USA), was dissolved in DMSO and used at 10 μM in keratinocytes.

### qRT-PCR analysis

RNA was harvested using the Qiagen RNeasy Mini Kit (Qiagen, USA) followed by iScript cDNA synthesis (Bio-Rad, USA). qRT-PCR was performed on a Roche LightCycler 480 II using the following parameters: 94°C for 10 seconds, 60°C for 10 seconds, 72°C for 10 seconds, for a total of 45 cycles^5^. All the expression data was normalized to the housekeeping gene, *L32*. Primer sequences are listed in Table 1.

**Table 1:**
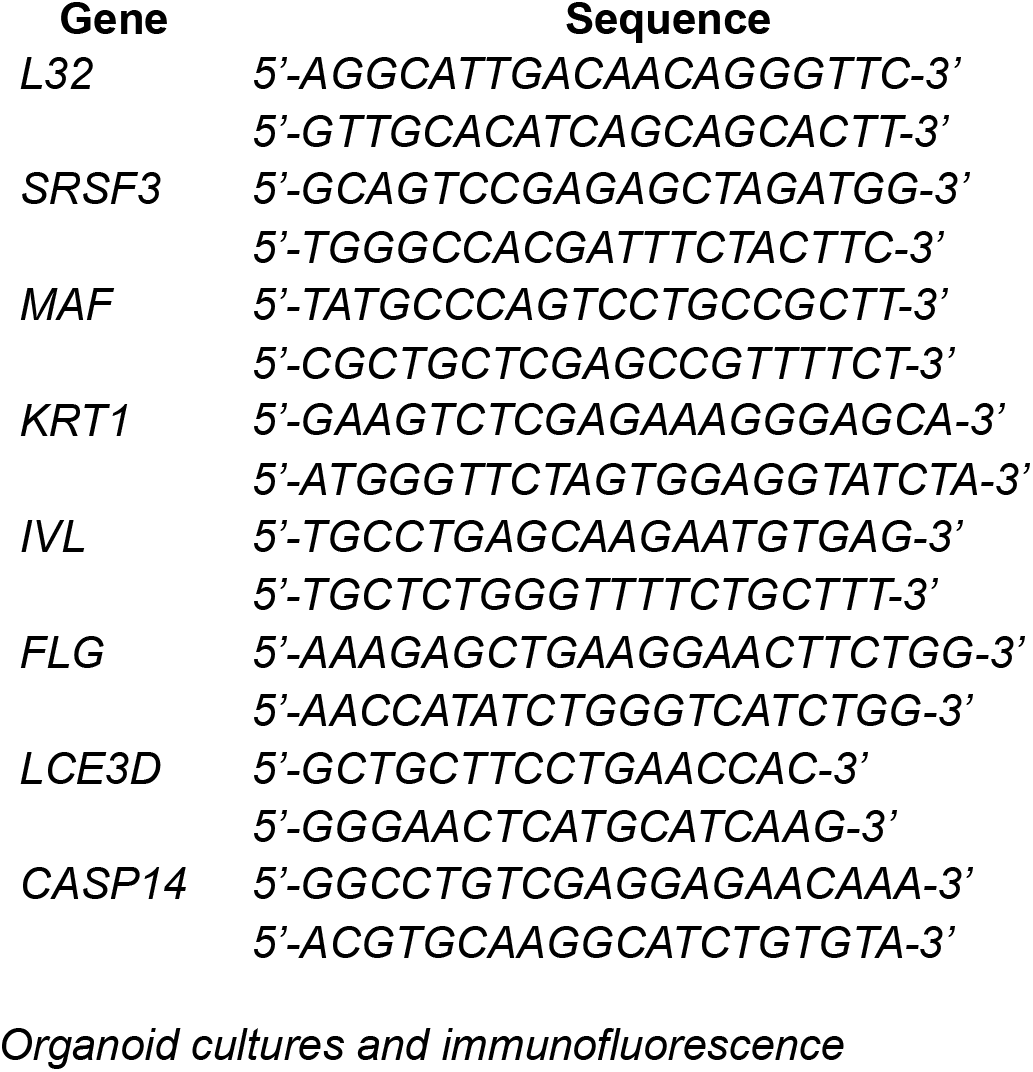
qRT-PCR primer sequences Gene Sequence.

### Organoid cultures and immunofluorescence

Primary keratinocytes were seeded onto human devitalized dermis^5^ and cultured at the air-liquid interface in the presence or absence of SFI003 (10 μM) for seven days. The skin tissue was embedded in Optimal Cutting Temperature compound (Sakura Fineteck, USA) on day seven and sectioned onto polylysine slides (ThermoFisher Scientific, USA) for immunofluorescence staining. Tissues were fixed in methanol or acetone as previously described^5^ and stained with differentiation and proliferation markers along with Collagen VII and Hoechst 33342 nuclear stain (Table 2). Fluorescent signals were observed using a ZEISS AXIO Observer Z1 microscope (ZEISS, USA).

**Table 2:** Antibody specifications.

| Protein | Vendor | Dilution |
| --- | --- | --- |
| Hoechst 33342 | ThermoFisher Scientific | 1:5000 |
| Collagen VII | MilliporeSigma | 1:200 |
| Keratin 1 | BioLegend | 1:2000 |
| Loricrin | Abcam | 1:500 |
| Filaggrin | Abcam | 1:500 |
| Proliferation marker Ki-67 | ThermoFisher Scientific | 1:100 |

### Electronic medical records analysis

National electronic health records (EHR) from 2010 to present day were examined by Atropos Health, which includes encounters from both community hospitals and large provider practices in the contiguous U.S. states in a variety of care settings. The incidence of cSCC in patients with a history of atrial fibrillation or heart failure who received the SRSF3 inhibitor digoxin (n=4,149) or amiodarone (n=9,579) were compared to those who never received either drug but were on beta blockers (n=100,000; down sampled), a standard treatment for these cardiac conditions. The index date was defined as the first receipt of medication and patients with (1) a history of atrial fibrillation or heart failure at baseline, (2) no diagnosis of cSCC in the 3 months prior to or after study inclusion, and (3) no diagnosis of metastatic cancer at baseline were followed from index to diagnosis of cSCC at any time after 3 months post-inclusion. Treatment populations were propensity scored matched for age, sex, race, and various comorbidities using the formula log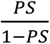, where PS is the probability of treatment. The propensity score was balanced amongst the treatment and control groups with a caliper of 0.25 to reduce confounding effects. The hazard ratios with 95% confidence intervals were calculated for the risk of cSCC incidence, and two-sided p-values of 0.05 were considered statistically significant.

## Results

Keratinocytes normally undergo terminal differentiation, present morphological differences, and lose proliferative capacity in the stratum spinosum, granulosum, and corneum layers of the epidermis. To demonstrate a functional role for SRSF3 in epidermal differentiation, we elected to use pharmacologic inhibition over genetic approaches, as homozygous deletion of SRSF3 causes embryonic lethality^3^. Using SFI003, a small molecule that degrades the SRSF3 protein via post-translational neddylation^6^, we reduced SRSF3 in primary human keratinocytes to approximately 50% of control levels without triggering visible cytotoxicity (Figure 1a). Depletion of SRSF3 for 48 hours induced the expression of both early (*MAF, KRT1*) and late (*IVL, FLG, LCE3D, CASP14*) markers of epidermal differentiation in subconfluent keratinocyte cultures that normally retain progenitor cells in the undifferentiated state (Figure 1a). We next used human skin organoids to assess the impact of SRSF3 inhibition on differentiation in three-dimensional epidermal tissue. Immunofluorescence analysis of these regenerated skin tissues further confirmed an upregulation of differentiation proteins. When compared to controls, SRSF3-inhibited tissues demonstrated KRT1 expression in the basal layer and stronger expression of loricrin (LOR) and filaggrin (FLG). Proliferation marker protein Ki-67 displayed a similar signal to control levels and epidermal stratification was comparable between treatment groups (Figure 1b). These findings suggest that SRSF3 is a negative regulator of differentiation in the skin.

**Fig 1.**
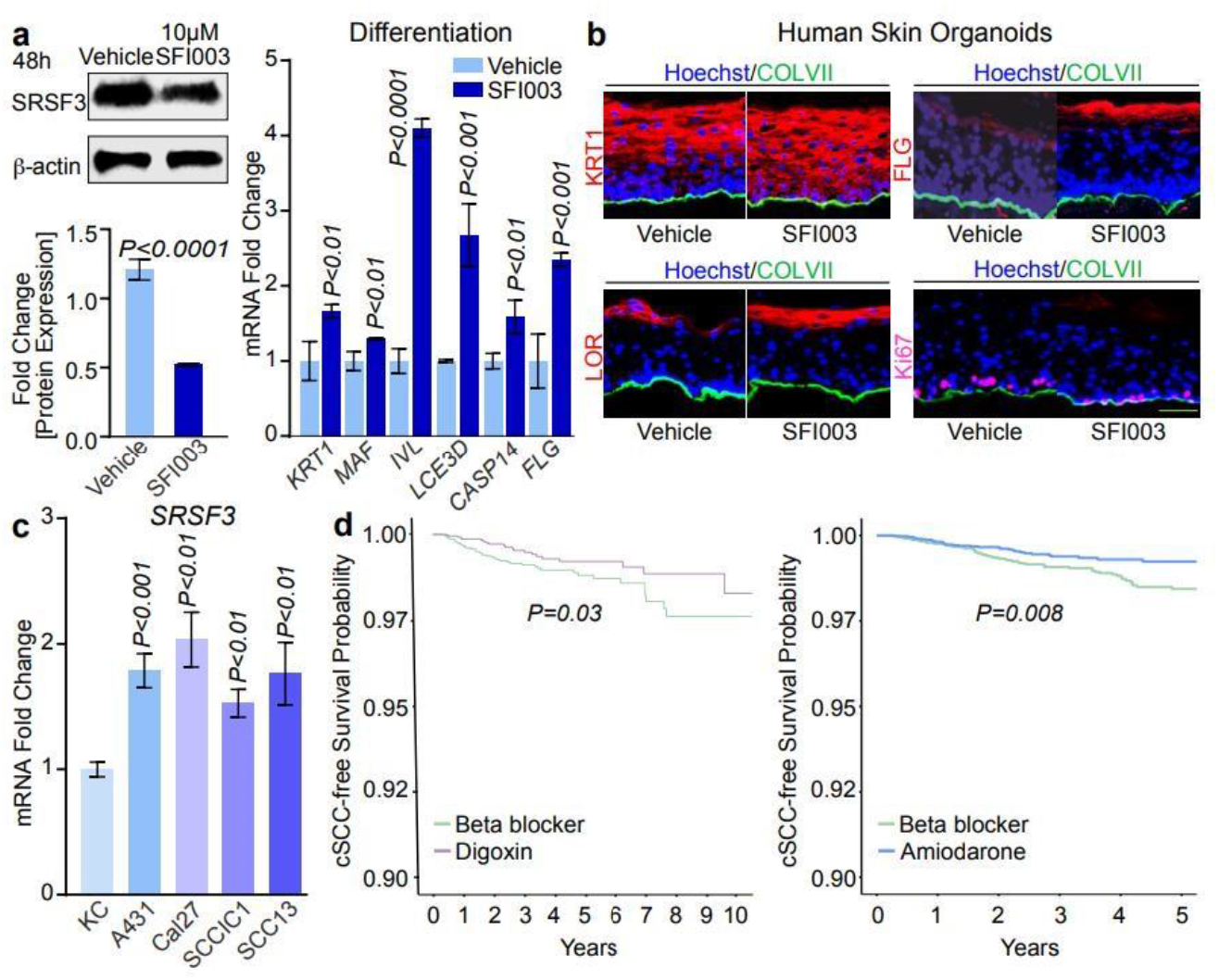
SRSF3 inhibition induces differentiation and is associated with increased cSCC-free survival. (a) Reduction of SRSF3 in primary human keratinocytes after 48 hours of SFI003 treatment (10 μM) with densitometry included (left). Differentiation gene expression in vehicle and SFI003-treated progenitor keratinocytes by qRT-PCR (right). (b) Immunofluorescence analysis of SRSF3-inhibited human skin organoids. Scale bar, 100 μm. (c) Quantitation of SRSF3 in cancer cell lines by qRT-PCR compared to primary keratinocytes. (d) Propensity score-matched cSCC-free survival probability of patients taking digoxin or amiodarone compared to beta blockers. P-values in (a) and (c) were calculated using a two-tailed t-test.

Dysregulated differentiation is an indicator of cSCC, as driver gene mutations disrupt normal differentiation pathways^7,8^. Our analysis of harmonized RNA-seq data capturing different clinicopathological stages of cSCC progression revealed that SRSF3 is upregulated in cSCC as well as precancerous actinic keratoses and decreased in epidermal differentiation^9-11^.

Interestingly, comparable levels of SRSF3 were detected in normal skin and keratoacanthomas, a self-healing form of cSCC that resolves through induction of differentiation^9^. These data are consistent with our observation of increased differentiation in both progenitor keratinocytes and skin organoids following pharmacologic inhibition of SRSF3 and suggest that targeting SRSF3 upregulation in cSCC might encourage malignant keratinocytes to terminally differentiate. We next examined the expression of SRSF3 in multiple squamous cancer cell lines and observed up to a two-fold increase in A431, CAL27, SCCIC1, and SCC13 cells compared to primary human keratinocytes (Figure 1c). This upregulation of the robustly expressed SRSF3 in cancer could contribute to the inability of these cell lines to terminally differentiate. To investigate whether inhibition of SRSF3 might reduce cSCC development, we leveraged the ability of digoxin and amiodarone, two medications commonly used to treat heart failure and arrhythmia, to reduce SRSF3 expression^12,13^. Using anonymized U.S. EHR data, we performed a retrospective cohort study comparing patients who received either digoxin or amiodarone to those who were given beta blockers, a standard treatment for heart failure and arrhythmia. After propensity score matching to adjust for potential confounders, both digoxin and amiodarone treatment were associated with a lower incidence of cSCC, as evidenced by higher cSCC-free survival compared to beta blockers (Figure 1d). Our analysis also identified a significant decrease in the hazard ratio (HR) of the digoxin (0.57, 95% CI 34-95%) and amiodarone (0.59, 95% CI 40-87%) groups compared to the control group. Our findings suggest that targeting SRSF3 with two distinct pharmacologic inhibitors reduces the risk of developing cSCC in humans.

## Discussion

In conclusion, our results demonstrate that SRSF3 limits differentiation in keratinocytes. Increased SRSF3 is associated with a poor prognosis in numerous cancers, including colorectal, prostate, and head and neck cancers^1^. Knockdown of SRSF3 in a colorectal cell line, HCT-116, led to suppressed xenograft growth in mice, while SFI003 displayed antitumor activity and curbed tumor growth in HCT-116 xenograft models^6^. SRSF3 genetic ablation or inhibition led to suppressed tumor growth in colorectal cancer models^6^, whereas its role in epidermal dedifferentiation to fuel cSCC is a novel concept. A potential limitation of this study is its focus on SRSF3 regulation of differentiation. We plan to explore other cancer phenotypes mediated by SRSF3, such as invasion and metastasis, beyond this pilot study. The downstream signaling pathways associated with SRSF3 have been explored in numerous cancers but require further examination in cSCC. In this study, we have established that targeting SRSF3 inhibition has therapeutic benefits for cSCC and can be further optimized as a promising treatment.

## References

1. Li D, Yu W, Lai M. Towards understandings of serine/arginine-rich splicing factors. Acta Pharm Sin B 2023; 13: 3181–207.

2. Jia R, Zheng ZM. Oncogenic SRSF3 in health and diseases. Int J Biol Sci 2023; 19: 3057–76.

3. Zhou Z, Gong Q, Lin Z et al. Emerging Roles of SRSF3 as a Therapeutic Target for Cancer. Front Oncol 2020; 10: 577636.

4. More DA, Kumar A. SRSF3: Newly discovered functions and roles in human health and diseases. Eur J Cell Biol 2020; 99: 151099.

5. Srivastava A, Tommasi C, Sessions D, Mah A, Bencomo T, Garcia JM, et al. MAB21L4 Deficiency Drives Squamous Cell Carcinoma via Activation of RET. Cancer Research. 2022 Sep 2;82(17):3143–57.

6. Zhang Y, Wang M, Meng F et al. A novel SRSF3 inhibitor, SFI003, exerts anticancer activity against colorectal cancer by modulating the SRSF3/DHCR24/ROS axis. Cell Death Discov 2022; 8: 238.

7. Bailey P, Ridgway RA, Cammareri P, Treanor-Taylor M, Bailey UM, Schoenherr C, et al. Driver gene combinations dictate cutaneous squamous cell carcinoma disease continuum progression. Nat Commun. 2023 Aug 25;14(1):5211.

8. Fania L, Didona D, Di Pietro FR et al. Cutaneous Squamous Cell Carcinoma: From Pathophysiology to Novel Therapeutic Approaches. Biomedicines 2021; 9.

9. Bencomo T, Lee CS. Gene expression landscape of cutaneous squamous cell carcinoma progression. Br J Dermatol 2024; 191: 760–74.

10. Kim, D.S., Risca, V.I., Reynolds, D.L. et al. The dynamic, combinatorial cis-regulatory lexicon of epidermal differentiation. Nat Genet 2021; 53, 1564–1576.

11. Takashima, S., Sun, W., Otten, A. B., Cai, P., Peng, S. I., Tong, E., Bui, J., Mai, M., Amarbayar, O., Cheng, B., Odango, R. J., Li, Z., Qu, K., & Sun, B. K. Alternative mRNA splicing events and regulators in epidermal differentiation. Cell Reports 2024; 43(3).

12. Lu GY, Liu ST, Huang SM et al. Multiple effects of digoxin on subsets of cancer-associated genes through the alternative splicing pathway. Biochimie 2014; 106: 131–9.

13. Chang YL, Liu ST, Wang YW et al. Amiodarone promotes cancer cell death through elevated truncated SRSF3 and downregulation of miR-224. Oncotarget 2018; 9:13390–406.

